# Dot6 is a major regulator of cell size and a transcriptional activator of ribosome biogenesis in the opportunistic yeast *Candida albicans*

**DOI:** 10.1101/401778

**Authors:** Julien Chaillot, Jaideep Malick, Adnane Sellam

## Abstract

In most species, size homeostasis appears to be exerted in late G1 phase as cells commit to division, called Start in yeast and the Restriction Point in metazoans. This size threshold couples cell growth to division and thereby establishes long-term size homeostasis. Our former investigations have shown that hundreds of genes markedly altered cell size under homeostatic growth conditions in the opportunistic yeast *Candida albicans*, but surprisingly only few of these overlapped with size control genes in the budding yeast *Saccharomyces cerevisiae*. Here, we investigated one of the divergent potent size regulators in *C. albicans*, the Myb-like HTH transcription factor Dot6. Our data demonstrated that Dot6 is a negative regulator of Start and also acts as a transcriptional activator of ribosome biogenesis (*Ribi*) genes. Genetic epistasis uncovered that Dot6 interacted with the master transcriptional regulator of the G1 machinery, SBF complex, but not with the *Ribi* and cell size regulators Sch9, Sfp1 and p38/Hog1. Dot6 was required for carbon-source modulation of cell size and it is regulated at the level of nuclear localization by TOR pathway. Our findings support a model where Dot6 acts as a hub that integrate directly growth cues via the TOR pathway to control the commitment to mitotic division at G1.

## Introduction

In a eukaryotic organism, cell size homeostasis is maintained through a balanced coordination between cell growth and division. In the last half century, a major focus of cell biology has been the study of cell division, but how eukaryotic cells couple growth to division to maintain a homeostatic size remains poorly understood. In most eukaryotic organisms, reaching a critical cell size appears to be crucial for commitment to cell division in late G1 phase, called Start in yeast and the Restriction Point in metazoans (Turner *et al.* 2012). Start is dynamically regulated by nutrient status, pheromone and stress, and facilitates adaptation to changing environmental conditions in microorganisms to maximize their fitness (Lenski and Travisano 1994; Kafri *et al.* 2016).

Different genome-wide genetic analyses have been accomplished in different model organisms to uncover the genetic determinism of Start and cell size control in eukaryotes. Screens of *Saccharomyces cerevisiae* mutants has identified many ribosome biogenesis (*Ribi*) genes as small size mutants (*whi*) (Jorgensen *et al.* 2002; Dungrawala *et al.* 2012; Soifer and Barkai 2014), and revealed two master regulators of *Ribi* gene expression, the transcription factor Sfp1 and the AGC family kinase Sch9, as the smallest mutants (Jorgensen *et al.* 2004). These observations lead to the hypothesis that the rate of ribosome biogenesis is a critical element of the metric that dictates cell size (Jorgensen *et al.* 2004; Schmoller and Skotheim 2015). Sfp1 and Sch9 are critical effectors of the TOR pathway and form part of a dynamic, nutrient-responsive network that controls the expression of *Ribi* genes and ribosomal protein genes (Jorgensen *et al.* 2004; Marion *et al.* 2004; Urban *et al.* 2007; Lempiainen *et al.* 2009). Sch9 is phosphorylated and activated by TOR, and in turn inactivates a cohort of transcriptional repressors of RP genes called Dot6, Tod6 and Stb3 (Huber *et al.* 2011).

*Candida albicans* is a diploid ascomycete yeast that is an important commensal and opportunistic pathogen in humans. While *C. albicans* and *S. cerevisiae* colonize different niches, common biological features are shared between the two yeasts including the morphological trait of budding, and core cell cycle and growth regulatory mechanisms (Berman 2006; Cote *et al.* 2009). *C. albicans* has served as an important evolutionary milestone with which to assess evolutionary conservation of biological mechanism, and recent evidence suggests a surprising extent of rewiring of central signalling, transcriptional and metabolic networks as compared to *S. cerevisiae* (Lavoie *et al.* 2009; Blankenship *et al.* 2010; Li and Johnson 2010; Sandai *et al.* 2012). To assess the conservation of the size control network, we performed recently a quantitative genome-wide analysis of a systematic collection of gene deletion strains in *C. albicans* (Sellam *et al.* 2016; Chaillot *et al.* 2017). Our screens uncovered that cell size in *C. albicans* is a complex trait that depends on diverse biological processes such as ribosome biogenesis, mitochondrial functions, cell cycle control and metabolism. In addition to conserved mechanisms and regulators previously identified in *S. cerevisiae* and metazoans, we uncovered many novel regulatory circuits that govern critical cell size at Start specifically in *C. albicans*. In particular, we delineate a novel stress-independent function of the p38/HOG MAPK pathway as a critical regulator of both growth, division, and poised to exert these functions in a nutrient-sensitive manner (Sellam *et al.* 2016). Interestingly, some of the size genes identified were required for fungal virulence, suggesting that cell size homeostasis may be elemental to *C. albicans* fitness inside the host.

An unexpectedly potent negative Start regulator that emerges from our systematic screen was *Dot6*, which encodes a Myb-like HTH transcription factor that binds to the PAC (Polymerase A and C) motif CGATG (Zhu *et al.* 2009; Sellam *et al.* 2016; Chaillot *et al.* 2017). *dot6* was among the smallest mutant identified by our screen. *C. albicans* Dot6 is the ortholog of two redundant transcriptional repressors of rRNA and *Ribi* gene expression called Dot6 and Tod6 in *S. cerevisiae*, which are antagonized by Sch9, and which cause only a minor large size phenotype when deleted together (Huber *et al.* 2011). Here, we show that the *C. albicans* Dot6 is a potent size regulator that govern critical cell size at Start and, in an opposite role than in *S. cerevisiae*, Dot6 acts as a transcriptional activator of *RiBi* genes. We also showed that the TOR pathway relays nutrient-dependent signal for size control to the Start machinery via Dot6. Genetic interactions with deletions of different known Start regulators revealed epistatic interaction with the master transcriptional regulator of the G1-S transition, SBF complex (Swi4-Swi6), but not with *SCH9, SFP1* or *HOG1*. These data emphasize the evolutionary divergence between *C. albicans* and *S. cerevisiae* and consolidate the role of Tor1-Dot6 network as a key cell size control mechanism in *C. albicans*.

## Materials and Methods

### Growth conditions and *C. albicans* Strains

The strains used in this study are listed in **Table S1**. *C. albicans* strains were generated and propagated using standard yeast genetics methods. For general propagation and maintenance conditions, the strains were cultured at 30°C in yeast-peptone-dextrose (YPD) medium supplemented with uridine (2% Bacto-peptone, 1% yeast extract, 2% dextrose, and 50 µg/ml uridine) or in Synthetic Complete medium (SC; 0.67% yeast nitrogen base with ammonium sulfate, 2% glucose, and 0.079% complete supplement mixture). The *DOT6*-△[1555-1803] truncated mutant was generated by inserting a STOP codon using CRISPR-Cas9 mutagenesis system (Vyas *et al.* 2015). gRNA was generated by annealing the Dot6-sgRNA-Top and Dot6-sgRNA-Bottom primers. Repair template was created using Dot6-STOP-Top and Dot6-STOP-Bottom primers (**Table S2**). The *C. albicans* SC5314 strain was co-transformed with the linearized plasmid pV1093 containing Dot6-gRNA with the repair template using lithium acetate transformation procedure and selected in Nourseothricin (Jena Bioscience). *DOT6* truncation was confirmed by sequencing.

### Cell size assessment

Cell size distributions were obtained using the Z2-Coulter Counter (Beckman). *C. albicans* cells were grown overnight in YPD at 30°C, diluted 1000-fold into fresh YPD or SC media and grown for 4 hours at 30°C to an early log phase density of 5×10^6^ ‒10^7^ cells/ml. A fraction of 100 µl of log phase culture was diluted in 10 ml of Isoton II electrolyte solution, sonicated three times for 10s and the distribution measured at least 3 times on a Z2-Coulter Counter. Size distributions were normalized to cell counts in each of 256 size bins and size reported as the peak median value for the distribution. Data analysis and clustering of size distributions were performed using custom R scripts that are available on request.

### Start characterization

The critical cell size at Start was determined by plotting budding index as a function of size in synchronous G1 phase fractions obtained using a JE-5.0 elutriation rotor with 40 ml chamber in a J6-Mi centrifuge (Beckman, Fullerton, CA) as described previously (Tyers *et al.* 1993). *C. albicans* G1 phase cells were released in fresh YPD medium and fractions were harvested at an interval of 10 min to monitor bud index. For the *dot6* mutant and the WT strains, additional size fractions were collected to assess transcript levels of the *RNR1, PCL2* and *ACT1* using qPCR (quantitative real time PCR) as cells progressed through G1 phase at progressively larger sizes.

### Growth assays

*C. albicans* cells were resuspended in fresh SC at an OD_600_ of 0.05. A total volume of 99 μl cells was added to each well of a flat-bottom 96-well plate in addition to 1 μl of the corresponding stock solution of either rapamycin or cycloheximide (Sigma). Growth assay curves were performed in triplicate in 96-well plate format using a Sunrise(tm) plate-reader (Tecan) at 30°C under constant agitation with OD_600_ readings taken every 10 min for 30h.

### Cellular localization of Dot6

A *DOT6*/*dot6* heterozygous strain was GFP-tagged *in vivo* at the C-terminal region with a GFP-Arg4 PCR product as previously described (Gola *et al.* 2003). Transformants were selected on SC minus Arginine plates, and correct integration of the GFP tag was checked by PCR and sequencing (**Table S2**). Live-cell microscopy of Dot6-GFP was performed with a Leica DMI6000B inverted confocal microscope (Leica) and a C9100-13 camera CCD camera (Hamamatsu). The effect of TOR activity on Dot6-GFP localization was assessed as following: cells grown on SC medium were exposed to rapamycin (100 ng/ml) for 60 min, washed once with PBS buffer and immediately visualized. *C. albicans* vacuoles were stained using the CellTracker Blue CMAC dye (ThermoFisher) following the manufacturer’s recommended procedure.

### Size genetic epistasis

*dot6* mutant was subjected to epistatic analysis with deletions of known Start regulators (Sellam *et al.* 2016) (**Table S1**). Gene deletion was performed as previously described (Gola *et al.* 2003). The complete set of primers used to generate deletion cassettes and to confirm gene deletions are listed in **Table S2**. Size distribution of at least, two independent double mutants were determined. Epistasis was only noted if size distributions of a single and double mutant overlapped.

### Microarray transcriptional profiling

Overnight cultures of *dot6* mutant and WT strains were diluted to an OD_600_ of 0.1 in 1 L fresh YPD-uridine medium, grown at 30°C to an OD_600_ of 0.8 and separated into size fractions by using the Beckman JE-5.0 elutriation system at 16°C. A total of 10^8^ unbudded G1 phase cells were harvested, released into fresh YPD medium and grown for 10 min prior to harvesting by centrifugation and stored at ‒80°C. Total RNA was extracted using an RNAeasy purification kit (Qiagen) and glass bead lysis in a Biospec Mini 24 bead-beater. Total RNA was eluted, assessed for integrity on an Agilent 2100 Bioanalyzer prior to cDNA labeling, microarray hybridization and analysis (Sellam *et al.* 2009). The GSEA Pre-Ranked tool (http://www.broadinstitute.org/gsea/) was used to determine statistical significance of correlations between the transcriptome of the *dot6* mutant with a ranked gene list or GO biological process terms as described by Sellam *et al.* (Sellam *et al.* 2014). Data were visualized using the Cytoscape (Saito *et al.* 2012) and EnrichmentMap plugin (Merico *et al.* 2010).

### Expression analysis by qPCR

For qPCR experiments, cell cultures and RNA extractions were performed as described for the microarray experiment. cDNA was synthesized from 1µg of total RNA using the SuperScipt III Reverse Transcription kit (ThermoFisher). The mixture was incubated at 25°C for 10 min, 37°C for 120 min and 85°C for 5 min. 2U/µl of RNAse H (NEB) was added to remove RNA and samples were incubated at 37°C for 20 min. qPCR was performed using an iQ5 96-well PCR system (BioRad) for 40 amplification cycles with QuantiTect SYBR Green PCR master mix (Qiagen). The reactions were incubated at 50°C for 2 min, 95°C for 2min and cycled 40 times at 95°C, 15 s; 56°C, 30 s; 72°C, 1 min. Fold-enrichment of each tested transcripts was estimated using the comparative △△Ct method as described by Guillemette *et al.* (Guillemette *et al.* 2004). To evaluate the gene expression level, the results were normalized using Ct values obtained from Actin (*ACT1*, C1_13700W_A). Primer sequences used for this analysis are summarized in Supplemental **Table S2**.

### Data Availability

Strains and plasmids are available upon request. Supplemental files contain three figures (Figure S1-S3) and four tables (Table S1-S4) and are available at FigShare (DOI: https://doi.org/10.6084/m9.figshare.7008170.v1).

## Results

### Dot6 is a negative regulator of START in *C. albicans*

We have previously shown that the transcription factor Dot6 was required for cell size control in *C. albicans* (Sellam *et al.* 2016). A *dot6* mutant had a median size that was 21% (41fL) smaller than its congenic parental (52fL) or the complemented strains (51fL) (**Figure 1A**). Inactivation of *DOT6* resulted in a delayed exit from the lag phase (1.5h delay as compared to the WT) (**Figure 1B**). However, *dot6* had a doubling time comparable to the WT and the complemented strains during the log phase suggesting that the size reduction of *dot6* is not a growth rate-associated phenotype (**Figure 1B**). To ascertain that this effect was mediated at Start, we evaluated two hallmarks of Start, namely bud emergence and the onset of SBF-dependent transcription as a function of cell size in synchronous G1 phase cells obtained by elutriation. As assessed by median size of cultures for which 90% of cells had a visible bud, the *dot6* mutant passed Start after growth to 26fL, whereas a parental WT control culture became 90% budded at a much larger size of 61fL (**Figure 1C**). Importantly, in the same experiment, the onset G1/S transcription was accelerated in the *dot6* strain as judged by the peak in expression of the two representative G1-transcripts, the ribonucleotide reductase large subunit, *RNR1* and the cyclin *PCL2* (**Figure 1D-E**). These results unequivocally demonstrated that Dot6 regulates the cell size threshold at Start.

**Figure 1.**
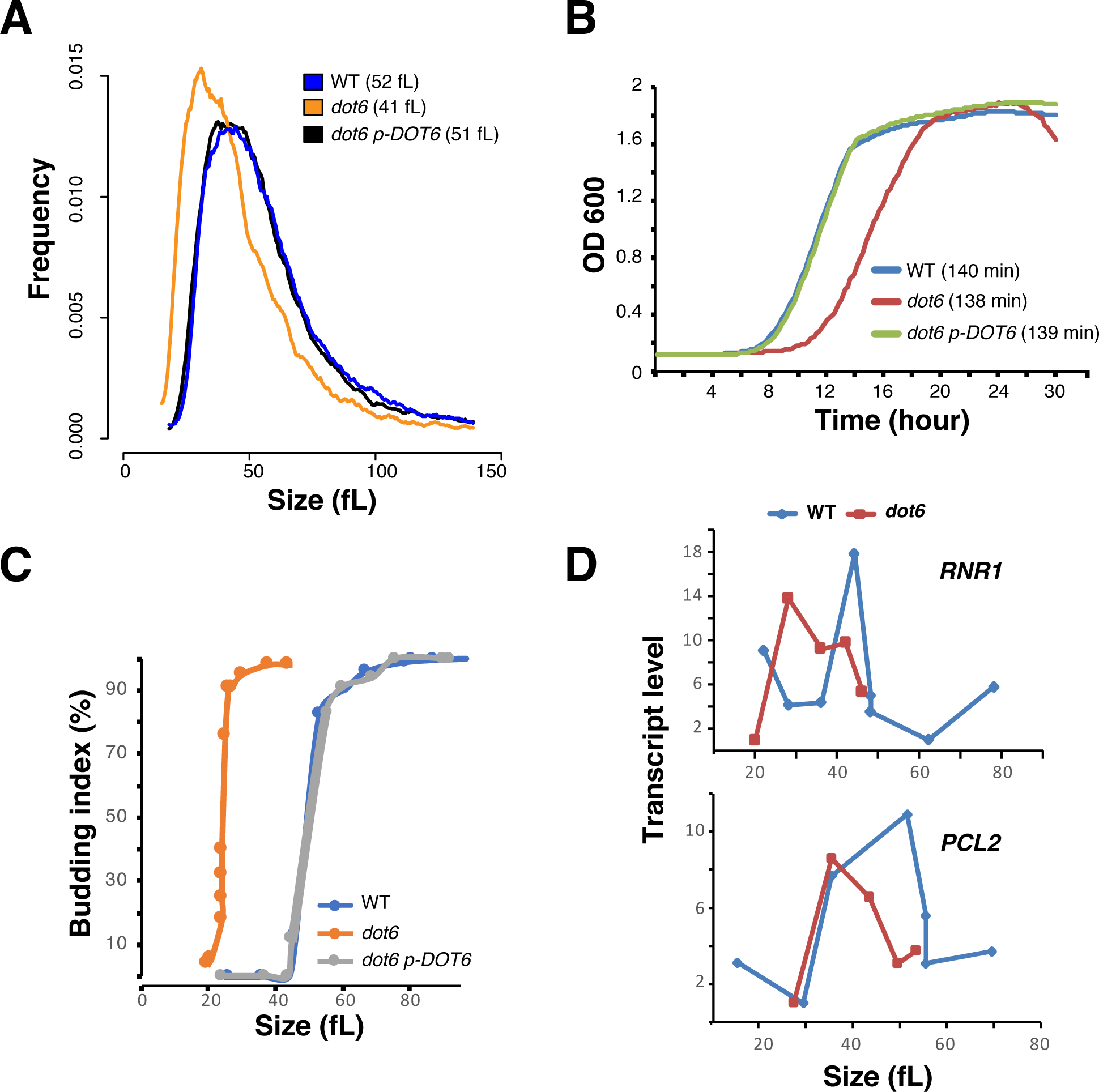
Dot6 is required for Start onset and cell size homeostasis. (A) Size distributions of the WT (SFY87), *dot6* mutant and the revertant strains. The median sizes of each strain are indicated in parentheses. (B) Growth of the WT (SFY87), *dot6* mutant and the revertant (*dot6* p-*DOT6*) strains in SC medium at 30°C. Doubling-times during the exponential phase of the growth for each strain are indicated in parentheses. (C-D) Start characterization of *dot6*. (C) Elutriated G1 phase daughter cells were released into fresh media and assessed for bud emergence as a function of size and G1/S transcription (D). *RNR1* and *PCL2* transcript levels were assessed by quantitative real-time PCR and normalized to *ACT1* levels.

### Dot6 interacts genetically with the SBF transcription factor complex

As cell size is a quantitative value, absolute changes in size between single and double mutants can be used to reveal genetic interactions between different genes to construct a cell size genetic interaction network (Jorgensen *et al.* 2002; Costanzo *et al.* 2004; De Bruin *et al.* 2004). To elucidate connections between Dot6 and previously identified Start regulators in *C. albicans* (Sellam *et al.* 2016), both *DOT6* alleles were deleted in different small size mutants including *hog1, sch9* and *sfp1* as well as the SBF large size mutant, *swi4*. Inactivating *DOT6* in either *sfp1, hog1* or *sch9* resulted in cells with smaller size as compared to their congenic strains suggesting that Dot6, Sfp1, Sch9 and the p38 kinase Hog1 act in different Start pathways (**Figure 2A-C**). Furthermore, inactivation of *DOT6* in the *swi4* mutant resulted in a large size comparable to that of *swi4* mutant indicating that Dot6 acts via SBF complex to control Start (**Figure 2D**). *SWI4* deletion is also epistatic to *DOT6* regarding the growth rate in liquid YPD medium confirming that both Dot6 and Swi4 act in a common pathway (**Figure 2E**). Given the absence of epistatic interaction between Dot6 and the known conserved *Ribi* and size regulators Sch9, Sfp1 and Hog1, our data uncovered a novel uncharacterized pathway that control the critical cell size threshold in *C. albicans* (**Figure 2F**).

**Figure 2.**
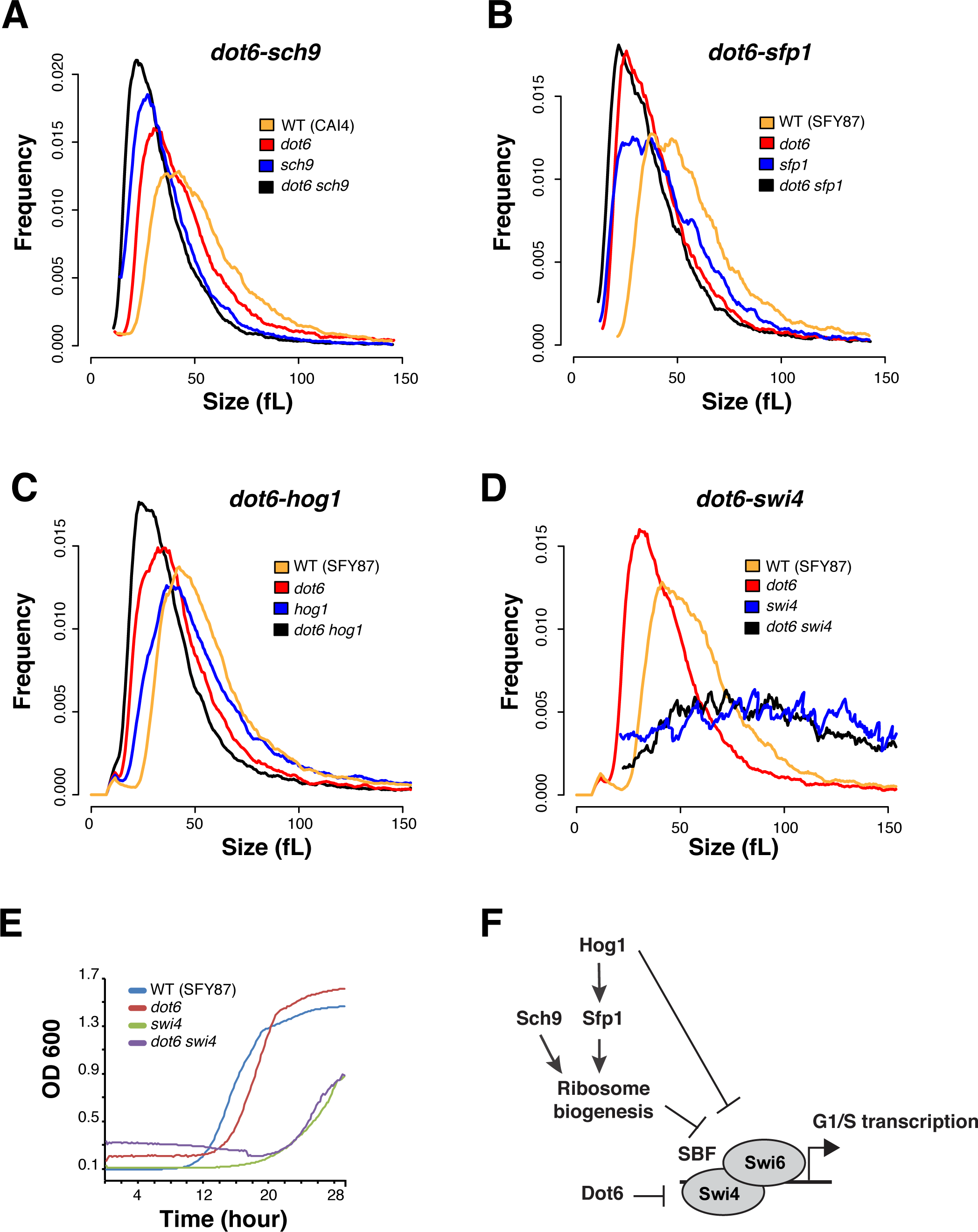
*DOT6* size epistasis. Evaluation of size epistasis between *dot6* and different potent Start mutations. *DOT6* was inactivated in *sch9* (A), *sfp1* (B), *hog1* (C) and *swi4* (D) mutants and the resulted double mutant strains were analyzed for cell size distribution. (E) *SWI4* deletion is epistatic to *DOT6* regarding the growth rate. Cells were grown in SC medium at 30°C under agitation with OD_600_ readings taken every 10 min for 30h. (F) Summary of *DOT6* genetic interactions with the *C. albicans* Start machinery.

### Dot6 is a positive regulator of ribosome biogenesis genes

Dot6 and its paralog Tod6 are both Myb-like transcription factors that repress *RiBi* genes in the budding yeast (Lippman and Broach 2009; Huber *et al.* 2011). To investigate the role of Dot6 in Start control in *C. albicans*, we performed genome-wide transcriptional profiling by microarray. G1-cells of both *dot6* mutants and the parental WT strain were collected by centrifugal elutriation and their transcriptomes were characterized. Gene Set Enrichment Analysis (GSEA) was used to correlate the *dot6* transcript profile with *C. albicans* genome annotations and gene lists from other transcriptional profiles experiments (Subramanian *et al.* 2005; Sellam *et al.* 2012) (**Table S2**). *dot6* mutant was unable to activate properly genes with functions mainly associated with protein translation, including ribosome biogenesis and structural constituents of the ribosome (**Figure 3A**). This suggest that in contrast to the role of its orthologue in *S. cerevisiae*, Dot6 in *C. albicans* is an activator of *RiBi*. Analysis of promoter region of transcript downregulated in *dot6* (transcript with 1.5-fold reduction using 5% FDR- **Table S3**) showed the occurrence of the PAC motif bound by Dot6 in all promoters of genes related to *RiBi* (**Figure 3B**). Furthermore, transcripts downregulated in *dot6* exhibited correlation with the set of genes repressed by the TOR complex inhibitor, rapamycin. This suggest that the evolutionary conserved *RiBi* transcription control by TOR is mediated fully or partially through Dot6. In support of the role of Dot6 in transcriptional control of *Ribi* genes and thus translation, *dot6* mutant exhibited an increased sensitivity to the protein translation inhibitor cycloheximide as compared to the WT and the revertant strains (**Figure 3C**).

**Figure 3.**
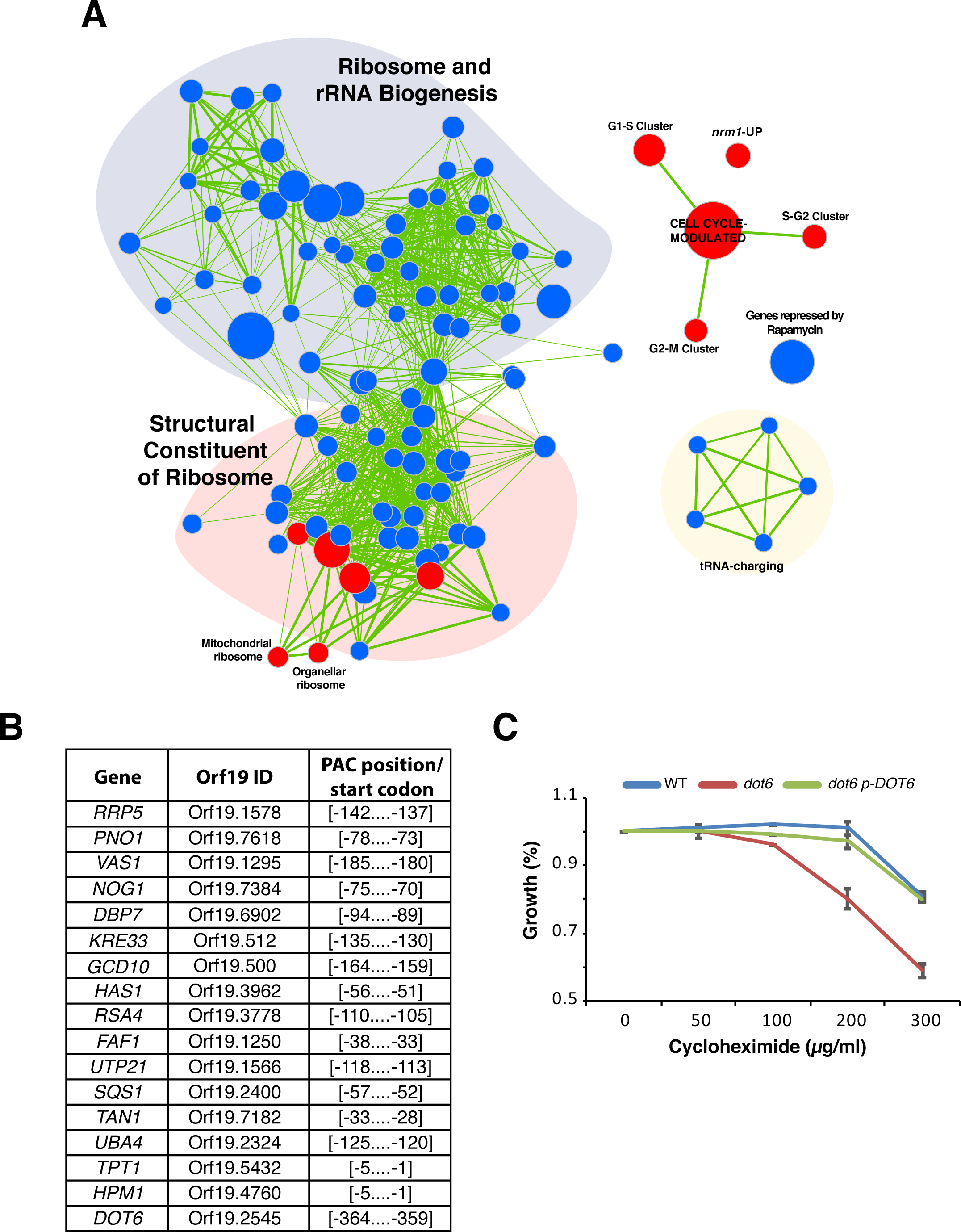
Dot6 is a positive regulator of ribosome biogenesis genes. (A) GSEA analysis of differentially expressed genes in a *dot6* mutant relative to the WT strain (SFY87). Cells were synchronized in G1 phase by centrifugal elutriation and released in fresh SC medium for 10 min and analyzed for gene expression profiles by DNA microarrays. Correlations of *dot6* up-regulated (red circles) and down-regulated (blue circles) transcripts are shown for biological processes, gene lists in different *C. albicans* mutants and experiments. The diameter of the circle reflects the number of modulated gene transcripts in each gene set. Known functional connections between related processes are indicated (green lines). Images were generated in Cytoscape with the Enrichment Map plug-in. (B) Occurrence of the PAC motif in the promoters of Dot6-modulated *Ribi* genes. The 400bp sequence upstream the start codon of downregulated genes in *dot6* (transcript with 1.5-fold reduction using 5% FDR) were scanned for the CGATG motif. (C) Effect of the translation inhibitor cycloheximide on the growth of the WT (SFY87), *dot6* mutant and the revertant (*dot6* p-*DOT6*) strains. Strains were grown on in SC medium at 30°C for 24 hours. Growth was calculated as percentage of OD_600_ of treated cells relatively to the non-treated controls. Results are the mean of three replicates.

The transcriptional programs characterizing the cell cycle G1/S transition in *C. albicans* (Cote *et al.* 2009) were hyperactivated in *dot6* mutant, which is a further support of the role of Dot6 as a negative regulator of G1/S transcription and Start (**Figure 3A**). Interestingly, *dot6* upregulated transcripts showed a significant correlation with those activated in the deletion mutant of the negative regulator of Start in *C. albicans*, Nrm1 (Ofir *et al.* 2012; Sellam *et al.* 2016).

### Dot6 localization is regulated by the TOR signalling pathway

TOR is a central signaling circuit that controls cellular growth in response to environmental nutrient status and stress in eukaryotes. In *S. cerevisiae*, the transcription factor Sfp1 and the AGC kinase Sch9 are critical effectors of the TOR pathway and form part of a dynamic, nutrient-responsive network that controls the expression of *Ribi* genes, ribosomal protein genes and cell size (Jorgensen *et al.* 2004; Urban *et al.* 2007; Lempiainen *et al.* 2009). In *S. cerevisiae*, both *sch9* and *sfp1* mutants are impervious to carbon source effects on Start (Jorgensen *et al.* 2004). In *C. albicans*, while *sfp1* and *sch9* mutants have the expected small size phenotype (Sellam *et al.* 2016), they still retain the ability to respond to carbon source shifts, unlike the *S. cerevisiae* counterparts (**Figure S1**). This suggest that the Sfp1-Sch9 regulatory circuit had rewired and is unlikely to rely on the nutrient status of the cell to Start control in *C. albicans*.

To assess whether the nutrient-sensitive TOR pathway communicates the nutrient status to Dot6, we first tested whether altering TOR activity by rapamycin could alter the subcellular localization of the Dot6-GFP fusion. In the absence of rapamycin, Dot6-GFP was localized exclusively in the nucleus in agreement with its role as a transcriptional activator under nutrient rich environment (**Figure 4A-C**). A weak GFP signal was also observed in the nucleolus and the vacuole. When cells were treated with rapamycin, Dot6-GFP was rapidly relocalized to the vacuole and only a small fraction remain in the nucleus (**Figure 4D-F**). The vacuolar localization of the Dot6-GFP was confirmed by its co localization with the CellTracker Blue-stained vacuoles (**Figure S2**). These data suggest that TOR pathway regulates the transcriptional function of Dot6.

**Figure 4.**
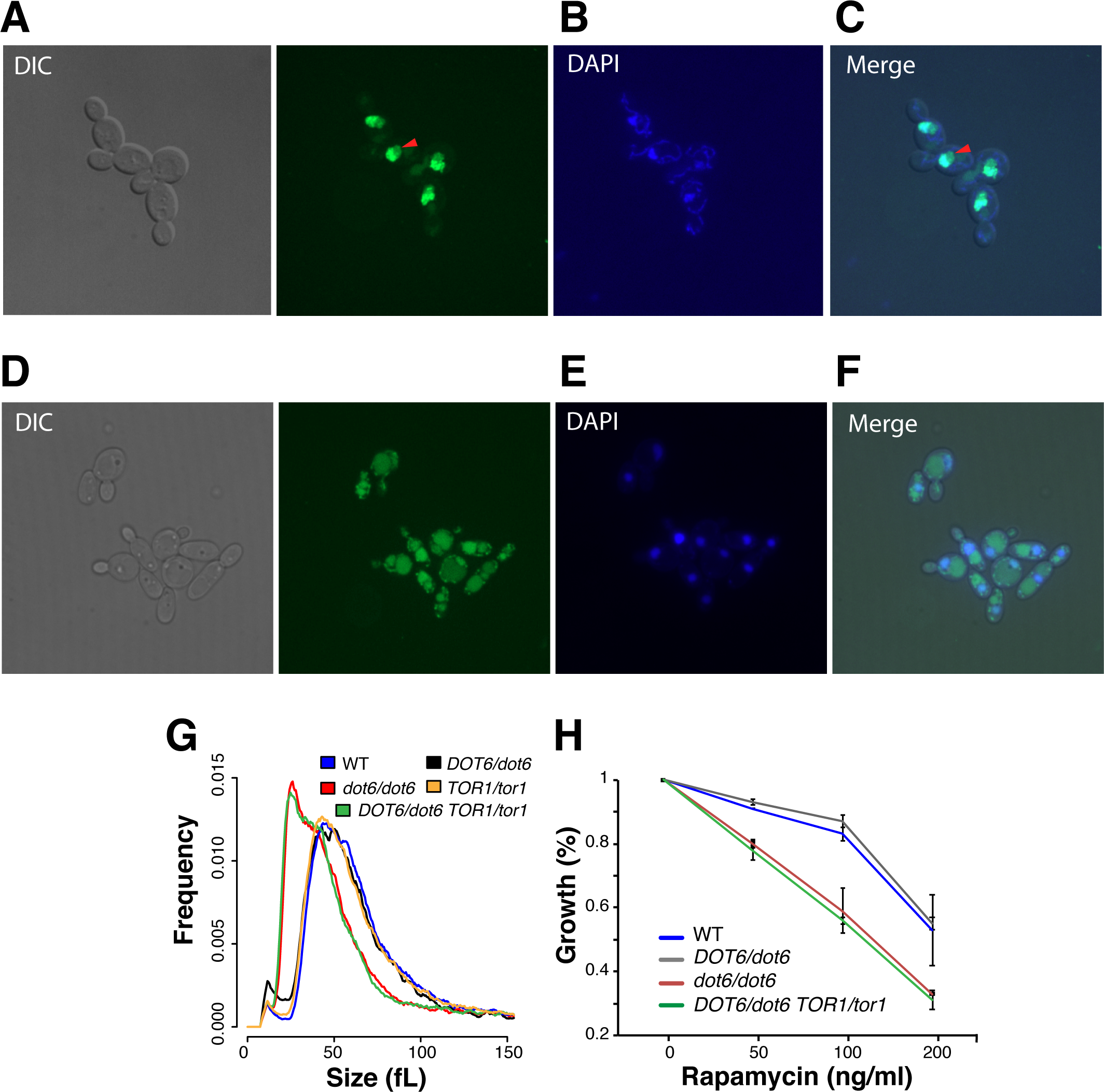
Dot6 localization is regulated by the TOR signalling pathway. (A-F) Dot6-GFP fluorescence was visualized using confocal microscopy in cells treated (D-F) or not (A-C) with the TOR pathway inhibitor, rapamycin. Exponentially grown cells in SC medium were treated with 100 ng/ml rapamycin for 1 hour. Nuclear and mitochondrial DNA were demarcated by DAPI staining (B and E). Red arrows indicate Dot6-GFP florescence in nucleolar regions. (G-H) *DOT6* and *TOR1* genetic interaction for cell size and growth in the presence of rapamycin based on complex haploinsufficiency concept. (G) Size distributions of the WT (SN250), the heterozygous (*DOT6*/*dot6*) and homozygous (*dot6*/*dot6*) *dot6* mutants, the heterozygous *TOR1*/*tor1* strain and the double heterozygous mutant *TOR1*/*tor1 DOT6*/*dot6.* (H) *DOT6* is epistatic to *TOR1* with respect to their sensitivity toward rapamycin. Strains were grown on in SC medium at 30°C for 24 hours. Growth was calculated as percentage of OD_600_ of rapamycin-treated cells relatively to the non-treated controls. Results are the mean of three replicates.

To assess whether the control of Dot6 activity by TOR impacts the cell size of *C. albicans*, we examined genetic interactions between *TOR1* and *DOT6* by size epistasis. As *TOR1* is an essential gene in *C. albicans*, we first tried to delete one allele in *dot6* homozygous mutant. However, all attempts to generate such mutant were unsuccessful suggesting a haplo-essentiality of *TOR1* in *dot6* mutant background. Subsequently, we analysed genetic interaction of *TOR1* and *DOT6* using complex haploinsufficiency (CHI) concept by deleting one allele of each gene and measured size distribution of the obtained mutant. While both *DOT6*/*dot6* and *TOR1*/*Tor1* mutants had no disenable size defect, the *TOR1*/*tor1 DOT6*/*dot6* strain exhibited cell size distribution similar to that of *dot6*/*dot6*, suggesting that *DOT6* is epistatic to *TOR1* (**Figure 4G**). Similarly, *DOT6* was also epistatic to *TOR1* with respect to their sensitivity toward rapamycin (**Figure 4H**). These data demonstrate that TOR pathway control cell size through Dot6.

### Dot6 is required for carbon-source modulation of cell size

The effect of different carbon sources was assessed on the size distribution of the *dot6* mutant and the WT. While the cell size of WT and the revertant strains was reduced by 12 % (47.6 ± 0.5 fL) when grown under the poor carbon source, glycerol, as compared to glucose (54.2 ± 0.5 fL), *dot6* size remain unchanged regardless the carbon source (**Figure 5A-B**). Similar finding was obtained when comparing cells growing on the non-fermentable carbon source, ethanol (data not shown). These results demonstrate that the transcription factor Dot6 is required for nutrient modulation of cell size. Furthermore, strain lacking *DOT6* was rate-limiting when grown in medium with glycerol as a sole source of carbon as compared to glucose (**Figure 5C**).

**Figure 5.**
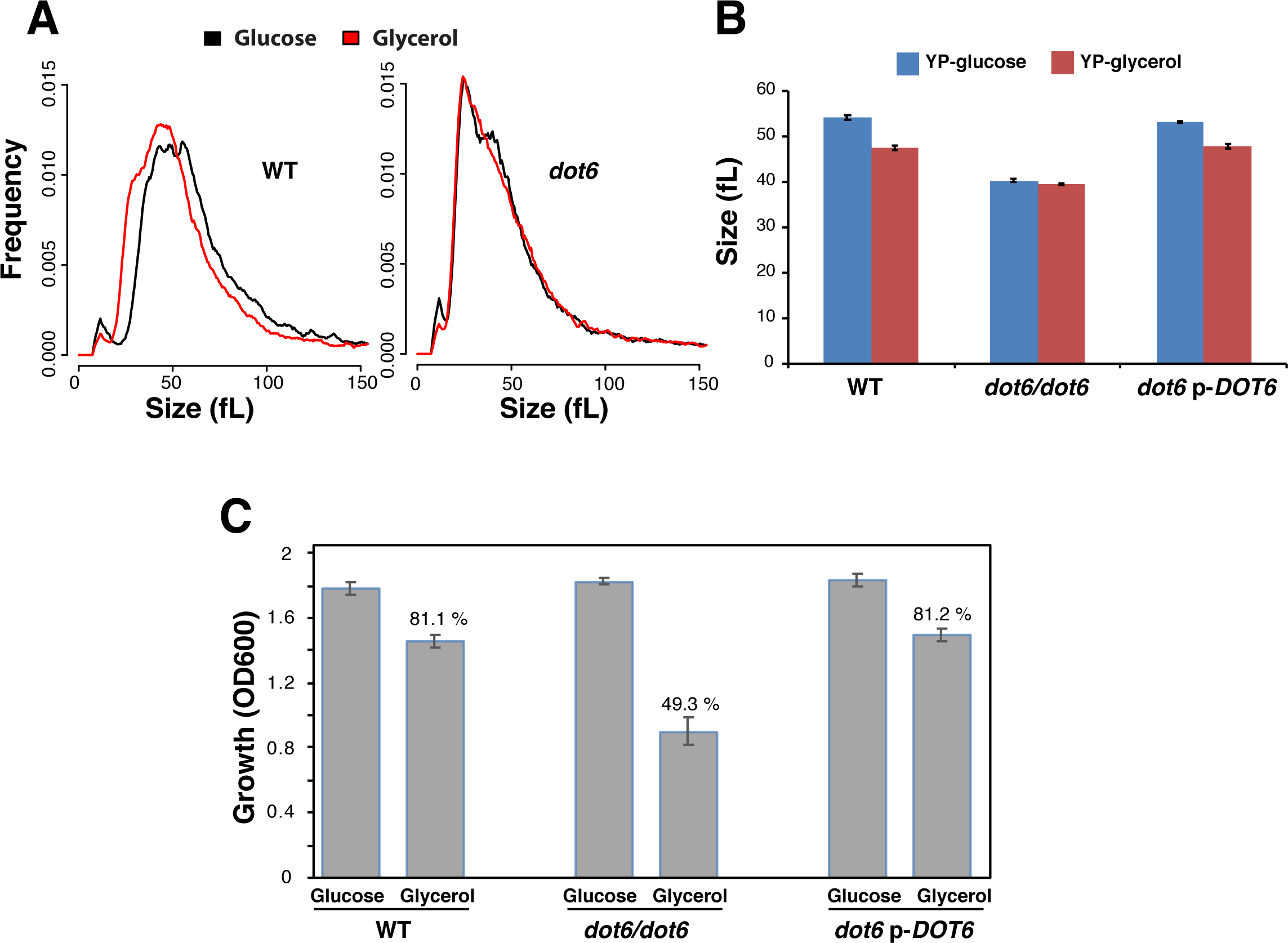
Dot6 is required for carbon-source modulation of cell size. (A) Cell size distribution of the WT and *dot6* mutant strains grown in medium with either glucose or glycerol as the sole source of carbon. (B) Median size of the WT (SFY87), *dot6* mutant and the revertant strains growing in synthetic glucose or glycerol medium. Results are the mean of three independent replicates. (C) Growth defect of *dot6* mutant in synthetic glycerol medium. The WT (SFY87), *dot6* mutant and the revertant strains were grown in synthetic glucose or glycerol medium at 30°C for 24 hours. Results are the mean of three independent replicates. The growth rate for each strain is indicated and represent the percentage of OD_600_ of cells grown in glycerol relatively to cells grown in glucose.

To check whether Dot6 localization is modulated by carbon sources, the subcellular localization of the Dot6-GFP fusion was tested in cells that grew in poor (glycerol) or in the absence of carbon sources. Neither the absence or the quality of carbon sources altered the nuclear localization of Dot6 (data not shown). This suggest that Dot6 govern the carbon-source modulation of cell size through a mechanism that is independent from its cellular relocalization

### The CTG-clade specific acidic domain of Dot6 is not required for size control

Our analysis unexpectedly reveals that Dot6 switched between activator and repressor transcriptional regulator of *Ribi* between *C. albicans* and *S. cerevisiae*, respectively. Sequence examination of *C. albicans* Dot6 protein revealed a C-terminal aspartate-rich domain that is similar to acidic activation domains of transcriptional activators. This Dot6 D-rich domain was found specifically in *C. albicans* and other related species of the CTG clade, and it was absent in Dot6 orthologs in *S. cerevisiae* and other ascomycetes (**Figure 6A**). To check whether the presence of this acidic domain corroborates with its function as transcriptional activator in *C. albicans*, we deleted this D-rich domain using CRISPR-Cas9 mutagenesis tool. Size distribution of the truncated *DOT6*-△[1555-1803] strain was indistinguishable from that of the WT parental strain (**Figure 6B**). The ability of *DOT6*-△[1555-1803] to activate two *Ribi* transcripts (*DBP7* and *KRE33*) was preserved which suggest that this domain is dispensable for the size control and gene expression activation functions of Dot6 (**Figure 6C**).

**Figure 6.**
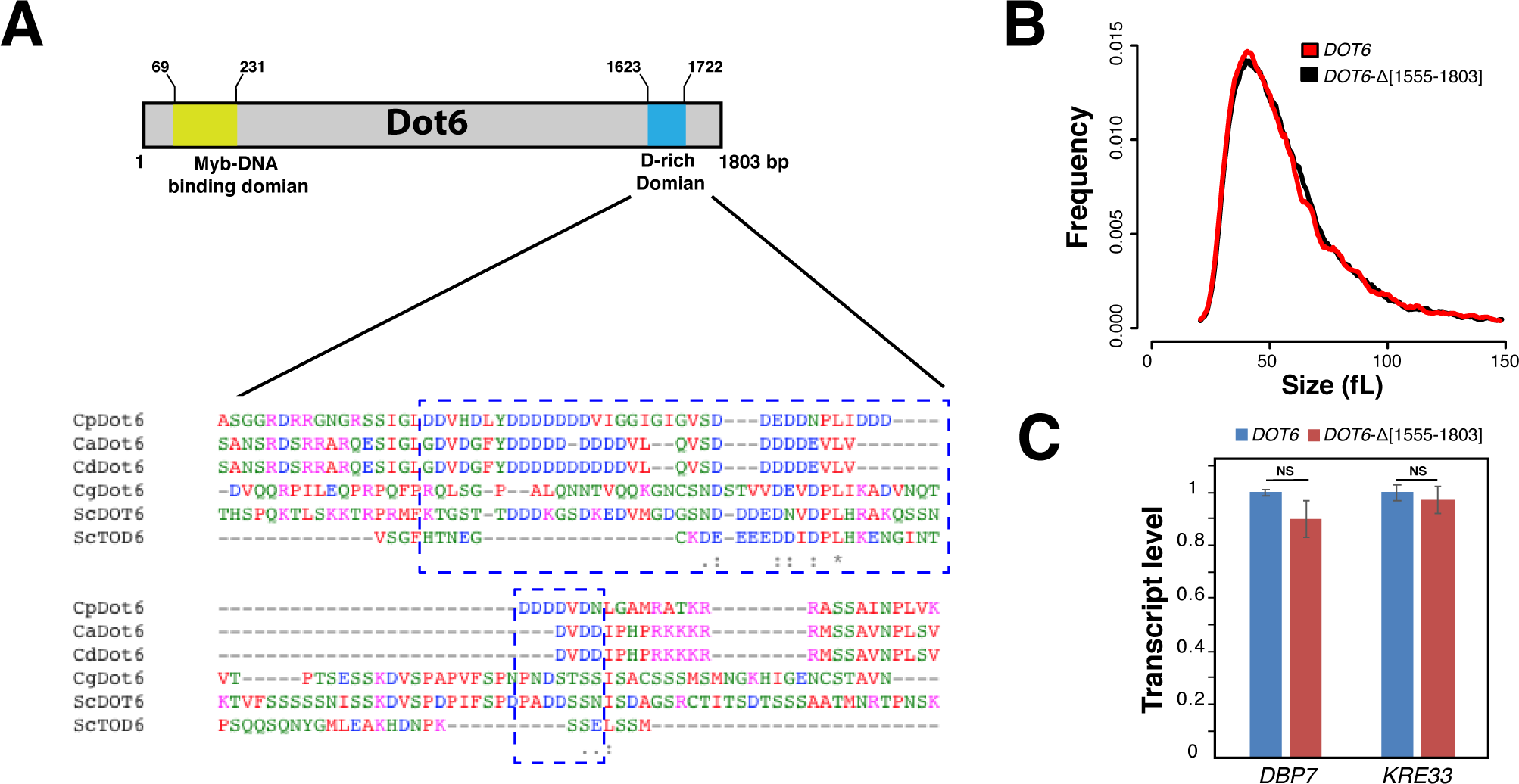
The CTG-clade specific acidic domain of Dot6 is not required for size control. (A) The C-terminal D-rich domain of Dot6 is conserved in the CTG clade species *C. albicans* (Ca), *C. parapsilosis* (Cp) and *C. dubliniensis* (Cd) but not in *S. cerevisiae* (Sc) and *C. glabrata* (Cg). Identical residues are indicated with asterisks. Conserved and semiconserved substitutions are denoted by colons and periods, respectively. (B) Cell size distribution of the WT (SC5314) and the truncated *DOT6*-△[1555-1803] strains. (C) Transcript levels of *Ribi* genes, including *DBP7* and *KRE33*, were evaluated in both WT (SC5314) and the truncated *DOT6*-△[1555-1803] strains. Transcript levels were calculated using the comparative CT method using the *ACT1* gene as a reference. Results are the mean of three replicates. For each transcript, fold changes in the WT and the truncated strains were not statistically significant (t-test). NS: not significant.

## Discussion

Although both *C. albicans* and *S. cerevisiae* share the core cell cycle and growth regulatory machineries, our previous investigations uncovered a limited overlap of the cell size phenome between the two fungi (Sellam *et al.* 2016; Chaillot *et al.* 2017). This finding is corroborated by recent evidences showing an extensive degree of rewiring and plasticity of both transcriptional regulatory circuits and signalling pathways across many cellular and metabolic processes between the two yeasts (Homann *et al.* 2009; Lavoie *et al.* 2009; Blankenship *et al.* 2010; Lavoie *et al.* 2010; Li and Johnson 2010; Childers *et al.* 2016). The plasticity of the global size network underscores the evolutionary impact of cell size as an adaptive mechanism to optimize fitness. Indeed, many size gene in *C. albicans* were linked to virulence which suggest that cell size is an important biological trait that contribute to the adaptation of fungal pathogens to their different niches (Sellam *et al.* 2016; Chaillot *et al.* 2017). So far, the requirement of Dot6 for the fitness of *C. albicans* inside its host was not tested yet, however, inactivation of *DOT6* led to the alteration of different virulence traits such as the sensitivity toward antifungals (Vandeputte *et al.* 2012). Moreover, while *dot6* mutant was able to form invasive filaments, the size of hyphae cells was significantly reduced which might impact the invasiveness of host tissues and organs (**Figure S3**). This reinforce the fact that control of cell size homeostasis is an important attribute for this *C. albicans* to persist inside its host.

We found that Dot6 is a major regulator of cell size in *C. albicans* as compared to *S. cerevisiae* emphasizing an evolutionary drift regarding the contribution of this transcription factor in size modulation. The potency of the *C. albicans* Dot6 in size control could be attributed to different facts. First, and in contrast to its role in *S. cerevisiae*, Dot6 is an activator of *Ribi* genes. This might explain the small size of *dot6* in *C. albicans* given the fact that inactivation of transcriptional activators of *Ribi* genes such as Sfp1 and Sch9 in either *S. cerevisiae* or *C. albicans* led to the acceleration of Start and, consequently, to a *whi* phenotype (Jorgensen *et al.* 2002; Dungrawala *et al.* 2012; Soifer and Barkai 2014; Sellam *et al.* 2016; Chaillot *et al.* 2017). Second, in *C. albicans,* Dot6 had an expanded genetic connectivity with both the critical SBF complex, that control the the G1/S transition, and also with the TOR growth and *Ribi* machineries, which might explain the influential role of Dot6 in size control.

Our findings support a model where Dot6 acts as a hub that might integrate directly growth cues via the TOR pathway to control the commitment to mitotic division at G1. This regulatory behavior is similar to the p38/HOG1 pathway that controls the *Ribi* regulon through the master transcriptional regulator, Sfp1, and acts upstream the SBF transcription factor complex to control division (Sellam *et al.* 2016). Meanwhile, our genetic interaction analysis showed that the *dot6 hog1* double mutant had an additive small size phenotype suggesting that both Dot6 and Hog1 act in parallel. This finding emphasizes that, in *C. albicans*, multiple signals are integrated at the level of G1 machinery to optimize adaptation to different conditions. Contrary to the p38/HOG pathway, Dot6 were required for both growth and size adjustment in response to glycerol suggesting that this transcription factor provides a nexus for integrating carbon nutrient status to the ribosome synthesis and Start machineries (**Figure 7**).

**Figure 7.**
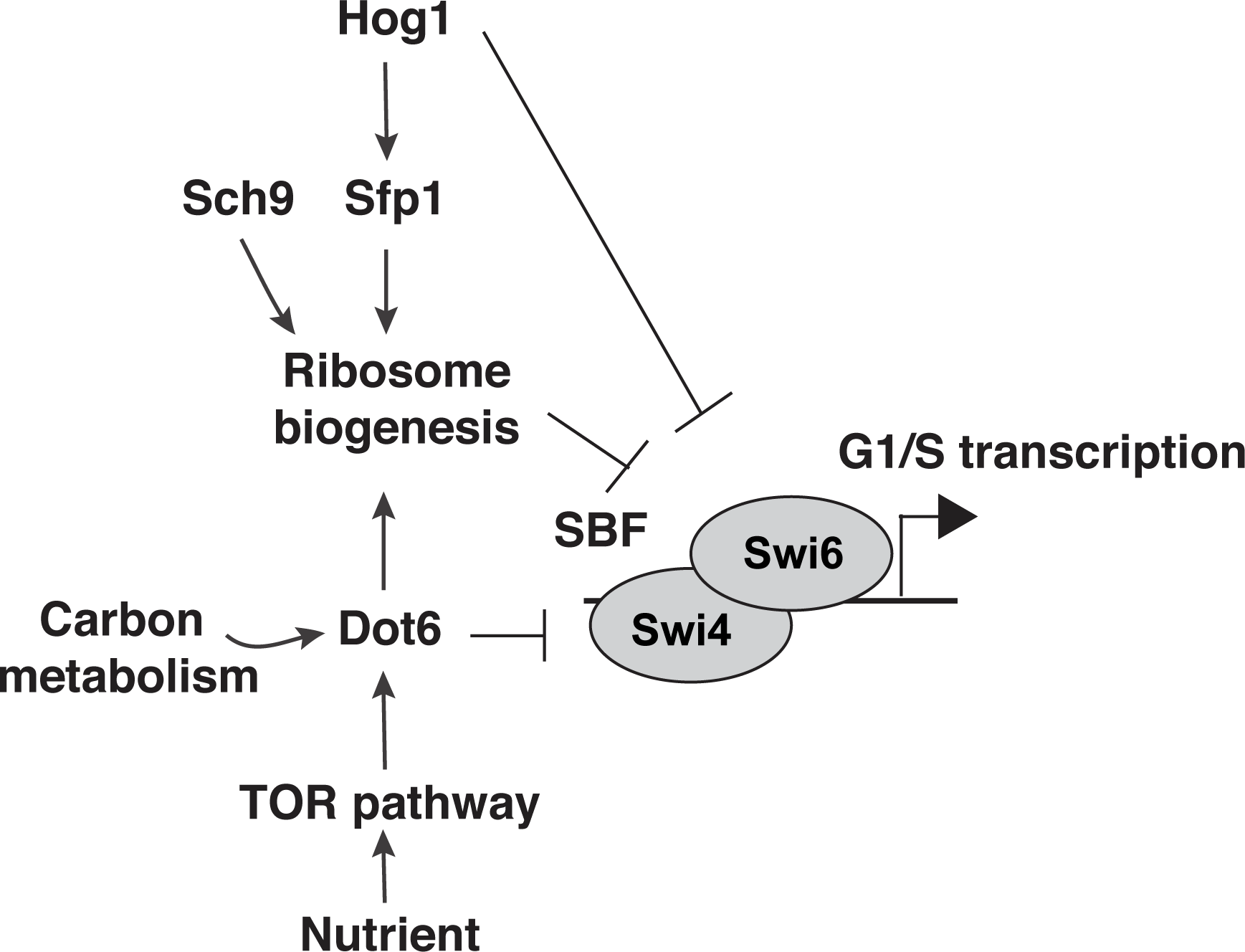
Schematic model of connections between Dot6 and Start control machinery in *C. albicans*.

## Supplementary Material

**Figure S1. A *sch9* mutant adjusts cell size in response to different carbon sources.**

Size distribution of log-phase cultures of the indicated WT (CAI4) and *sch9* strains grown in synthetic glucose (black curve) and glycerol (red) medium.

**Figure S2. Localization of Dot6-GFP in the vacuole when TOR pathway is compromised.**

Dot6-GFP fluorescence was visualized using confocal microscopy in cells treated with the TOR pathway inhibitor, rapamycin. Vacuoles were stained using CellTracker Blue CMAC dye. Red and blue arrows indicate Dot6-GFP florescence in the vacuole and the nucleus, respectively.

**Figure S3. Dot6 is required for size homeostasis of hyphal cells.**

Length of at least 20 hyphal cells of both WT (SFY87) and dot6 mutant grown on YPD supplemented with 10% fetal bovine serum (FBS) at 37°C. Bars represent the means ± standard errors of the means. *, *P* < 0.0003 using a two-tailed t-test.

**Table S1.** Strains used in the current study and their genotypes.

**Table S2**. Primer sequences used in the current study.

**Table S3**. Gene Set Enrichment Analysis (GSEA) of dot6 mutant transcriptome

**Table S4**. Transcripts differentially expressed in dot6 mutant using a 1.5-fold change cut-off and a 5% false discovery rate.

